# Identification of the glycosylphosphatidylinositol-specific phospholipase A2 (GPI-PLA2) of GPI fatty acid remodelling in *Trypanosoma brucei*

**DOI:** 10.1101/2023.04.18.536892

**Authors:** Zhe Ji, Rupa Nagar, Samuel M. Duncan, Maria Lucia Sampaio Guther, Michael A.J. Ferguson

**Affiliations:** Wellcome Centre for Anti-Infectives Research, School of Life Sciences, University of Dundee, Dundee DD1 5EH, United Kingdom

**Keywords:** Glycosylphosphatidylinositol, GPI, Trypanosoma brucei, phospholipase A2, PLA2, fatty acid remodelling

## Abstract

The biosynthesis of glycosylphosphatidylinositol (GPI) anchored proteins (GPI-APs) in the parasitic protozoan *Trypanosoma brucei* involves fatty acid remodelling of the GPI precursor molecules before they are transferred to protein in the endoplasmic reticulum. The genes encoding the requisite phospholipase A2 and A1 activities for this remodelling have thus far been elusive. Here, we identify a gene, Tb927.7.6110, that encodes a protein that is necessary and sufficient for GPI-phospholipase A2 (GPI-PLA2) activity in the procyclic form of the parasite. The predicted protein product belongs to the alkaline ceramidase, PAQR receptor, Per1, SID-1, and TMEM8 (CREST) superfamily of transmembrane hydrolase proteins and shows sequence similarity to Post-GPI-Attachment to Protein 6 (PGAP6), a GPI-PLA2 that acts after transfer of GPI precursors to protein in mammalian cells. The trypanosome Tb927.7.6110 GPI-PLA2 gene resides in a locus with two closely related genes Tb927.7.6150 and Tb927.7.6170, one of which (Tb927.7.6150) most likely encodes a catalytically inactive protein. The absence of GPI-PLA2 in the null mutant procyclic cells not only affected fatty acid remodelling but also reduced GPI anchor sidechain size on mature GPI-anchored procyclin glycoproteins. This reduction in GPI anchor sidechain size was reversed upon the add back of Tb927.7.6110 and of Tb927.7.6170, despite the latter not encoding GPI precursor GPI-PLA2 activity.

## Introduction

Glycosylphosphatidylinositol (GPI) anchored proteins (GPI-APs) are almost ubiquitous in the eukaryotes (1). The first complete GPI structure was determined for the variant surface glycoprotein (VSG) of the bloodstream form (BSF) of *Trypanosoma brucei*, the causative agent of human and animal African trypanosomiasis (2). Subsequent structures for rat Thy-1 antigen (3), human erythrocyte acetylcholinesterase (4), yeast Gas1p (5) and a myriad of other examples, reviewed in (1), have established the conserved and species/tissue-specific features of GPI anchors.

All GPI membrane anchors are based on a common underlying structure of ethanolamine-P-6Man*α*1-2Man*α*1-6Man*α*1-4GlcN*α*1-6PI (EtN-P-Man_3_GlcN-PI), where the amino group of the ethanolamine residue is in amide linkage to the *α*-carboxyl group of the C-terminal amino acid. This common core can then be decorated with carbohydrate sidechains and additional ethanolamine phosphate groups in a species- and tissue-specific manner (1). The phosphatidylinositol (PI) component can be diacyl-PI, a *lyso-*acyl-PI or an alkylacyl-PI (with or without an additional ester-linked fatty acid on the 2-position of the D-*myo*-inositol residue) and in some cases, like in yeast and some protozoa, an inositol-phospho-ceramide (1).

The biosynthesis of GPI anchors is quite well understood in *T. brucei* and other protozoan parasites, as well as in yeast and mammalian cells (1). As first noted in (6), the PI species of GPI anchors are different in lipid composition to the bulk PIs of their resident cells. The pathways in *T. brucei* and mammalian cells are summarised in (**Fig. 1**). Some of the key differences between GPI biosynthesis in *T. brucei* and mammalian cells lie in their inositol-acylation and lipid remodelling reactions. The latter are responsible for the aforementioned atypical PI compositions of GPI anchors.

**Figure 1:**
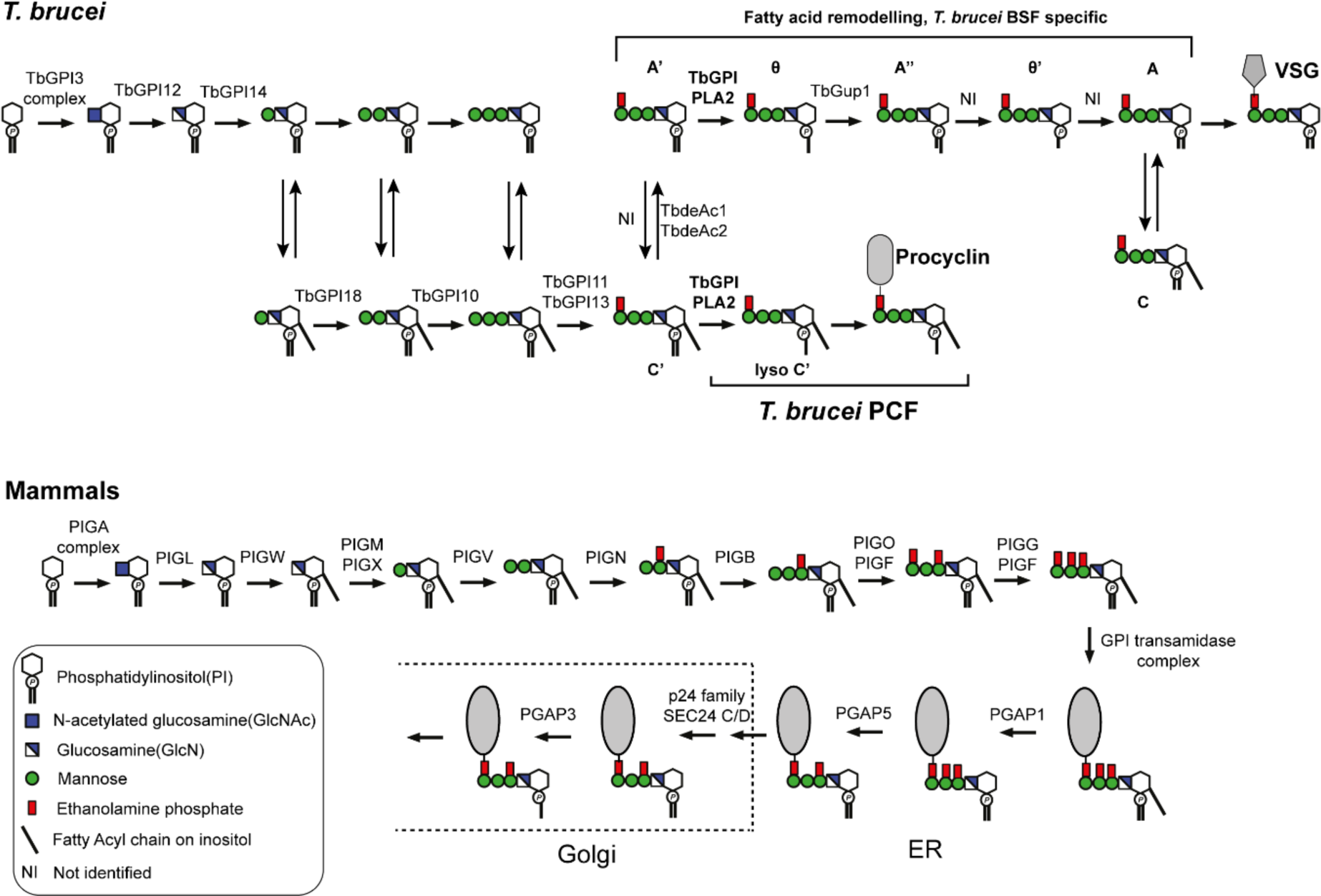
The GPI biosynthesis pathways in *T. brucei* and mammalian cells. The GPI biosynthesis pathway for *T. brucei* is shown for both the bloodstream form (BSF) and the procyclic form (PCF) of the parasite. After the addition of first mannose, all the GPI intermediates are in equilibrium between their inositol-acylated and non-acylated forms, as indicated by double arrows. The fatty acid remodelling occurs on the GPI intermediates before transferring to protein. By contrast, the GPI pathway for mammalian cells involves inositol-acylation at the level of GlcN-PI onwards, with removal of this fatty acid and lipid remodelling, by the PGAP enzymes, only after GPI transfer to protein.

With respect to inositol-acylation, in mammalian cells and yeast the 2-position of the inositol ring is acylated with predominantly palmitate (C16:0) in an acyltransferase reaction catalysed by PIG-W (GWT1 in yeast) using acyl-CoA as donor substrate and GlcN-PI as acceptor substrate to yield GlcN-(acyl)PI (7, 8). In *T. brucei*, there is no orthologue of PIG-W/GWT1 and the acyltransferase has not been identified. However, it is known that the donor substate is not acyl-CoA but most likely an endogenous ER membrane lipid (9, 10) and its earliest acceptor is Man_1_GlcN-PI, producing Man_1_GlcN-(acyl)PI (11). Further, the fatty acid transferred is more heterogeneous in *T. brucei* (a mixture of C14:0, C16:0, C18:0, C18:1 and C18:2) (12, 13). In mammalian cells and BSF *T. brucei*, the inositol-acyl chain can be removed by an inositol-deacylase (PGAP1 or deAc2) but this occurs after (14) or before (9–11) transfer of the GPI precursor to protein, respectively. The expression of deAc2 is tightly regulated and does not occur in PCF trypanosomes (15).

Lipid remodelling of GPIs was first described in *T. brucei* (16). In BSF cells, the diacyl-PI moiety of EtN-P-Man_3_GlcN-PI (glycolipid A’) is sequentially acted upon by: (i) an unidentified GPI-PLA2 to produce glycolipid *θ*, (ii) a myristoyltransferase (TbGup1) to produce glycolipid A’’(17), (iii) an unidentified GPI-PLA1 to produce glycolipid *θ*’ and (iv) an unidentified myristoyltransferase to yield the mature GPI precursor (glycolipid A) bearing a dimyristoyl-PI lipid that is competent for transfer to protein. In PCF cells, inositol de-acylation does not occur and fatty acid remodelling only proceeds as far as the action of GPI-PLA2 (18, 19).

In mammalian cells, lipid remodelling starts in the ER through the selection of *sn*-1-alkyl-2-acyl-(acyl)PI forms of GlcN-(acyl)PI (20). The unsaturated *sn*-2-acyl cain is then exchanged for a saturated (predominantly C18:0) fatty acid through the sequential action of a GPI-PLA2 (called PGAP3) (21) and an acyltransferase reaction dependent on PGAP2 (22). The resulting, generally fully saturated, alkylacyl-PI GPI anchor then associates with liquid ordered membrane domains (known as lipid rafts) for forward transport to the plasma membrane (21).

The mammalian fatty acid remodelling GPI-PLA2 enzyme (PGAP3) belongs to the (alkaline ceramidase, PAQR receptor, Per1, SID-1, and TMEM8 (CREST) superfamily of proteins (23). Another CREST superfamily member, TMEM8A/PGAP6, was also shown to be a GPI-PLA2 acting on certain GPI-APs to facilitate their release from mammalian cells in a soluble form (24, 25). The CREST superfamily includes several lipases that share a core structure of seven predicted transmembrane domains and five conserved residues (three histidine, one aspartic acid and one serine) (23).

In this paper, we identify three CREST superfamily members in *T. brucei* that are in tandem array on chromosome 7. We provide direct evidence that one of these genes encodes the hitherto elusive GPI-PLA2 of *T. brucei* GPI fatty acid remodelling, and we speculate on the possible functions of the other two.

## Results

### Identification of putative TbGPI-PLA2 genes

While the lipid remodelling reactions in GPI biosynthesis are different between *T. brucei* and mammalian cells (1), as indicated in (**Fig. 1**), both processes include GPI-specific phospholipase A2 reactions. We reasoned that the *T. brucei* GPI-PLA2(s) might have amino acid sequence similarity to PGAP3 and/or PGAP6, the two known mammalian GPI-PLA2s.

Using Delta BLASTp searches (26) we found three *T. brucei* genes (Tb927.7.6110, Tb927.7.6150 and Tb927.7.6170) in tandem array with 11-16% sequence identity to PGAP6 and seven predicted closely spaced putative transmembrane domains (**Fig. 2*A***). The predicted amino acid sequences of Tb927.7.6110, Tb927.7.6150 and Tb927.7.6170 have 50-66 % sequence identity and Tb927.7.6110 and Tb927.7.6170 possess all five of the conserved residues (3 His, 1 Ser and 1 Asp) of the CREST superfamily lipases, including PGAP6 (**Fig. 2*B***).

**Figure 2:**
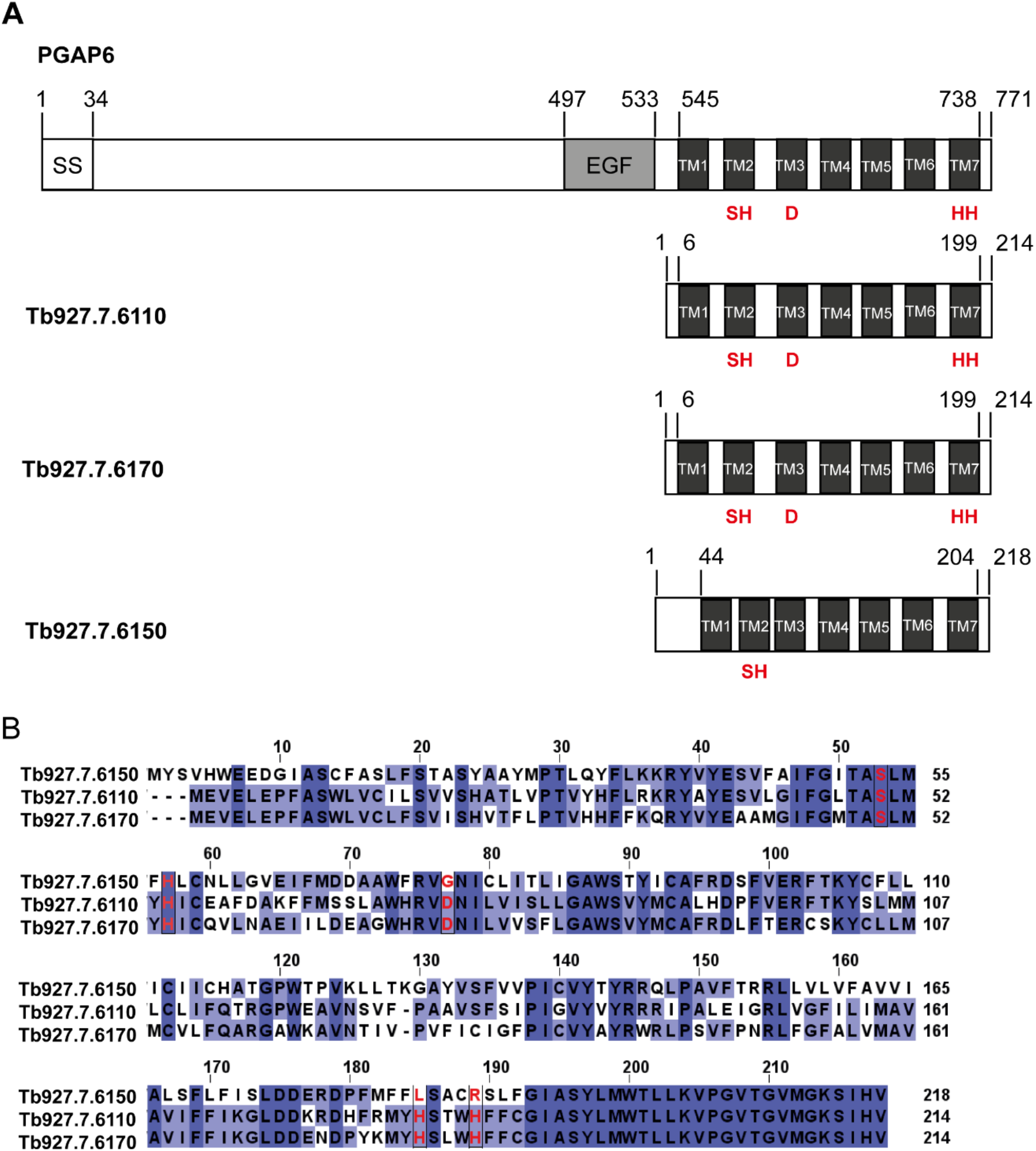
Similarity of PGAP6 to three predicted *T. brucei* gene products. The domain organisation of PGAP6 and the three related *T. brucei* genes are shown. PGAP6 contains an N-terminal signal sequence (SS) and an EGF-like domain (EGF) in front of its seven putative transmembrane domains (TMs). The TM domains of PGAP6, Tb927.7.6110 and Tb927.7.6170 share all five conserved residues of the CREST lipase family (in red), whereas Tb927.7.6150 shares only two of these conserved residues. (**B**) shows the sequence alignment of the three *T. brucei* gene products (Tb927.7.6110, Tb927.7.6150 and Tb927.7.6170).

### Tb927.7.6110 is necessary and sufficient for TbGPI-PLA2 activity

Since GPI biosynthesis is not essential to PCF cells (27), we made a Tb927.7.6110/6150/6170 locus null mutant by gene replacement, as indicated in (**Fig. 3*A***). Confirmatory Southern blot data are in (**Fig. S1**). The resulting null mutant was viable with similar growth kinetics to wild-type cells.

**Figure 3:**
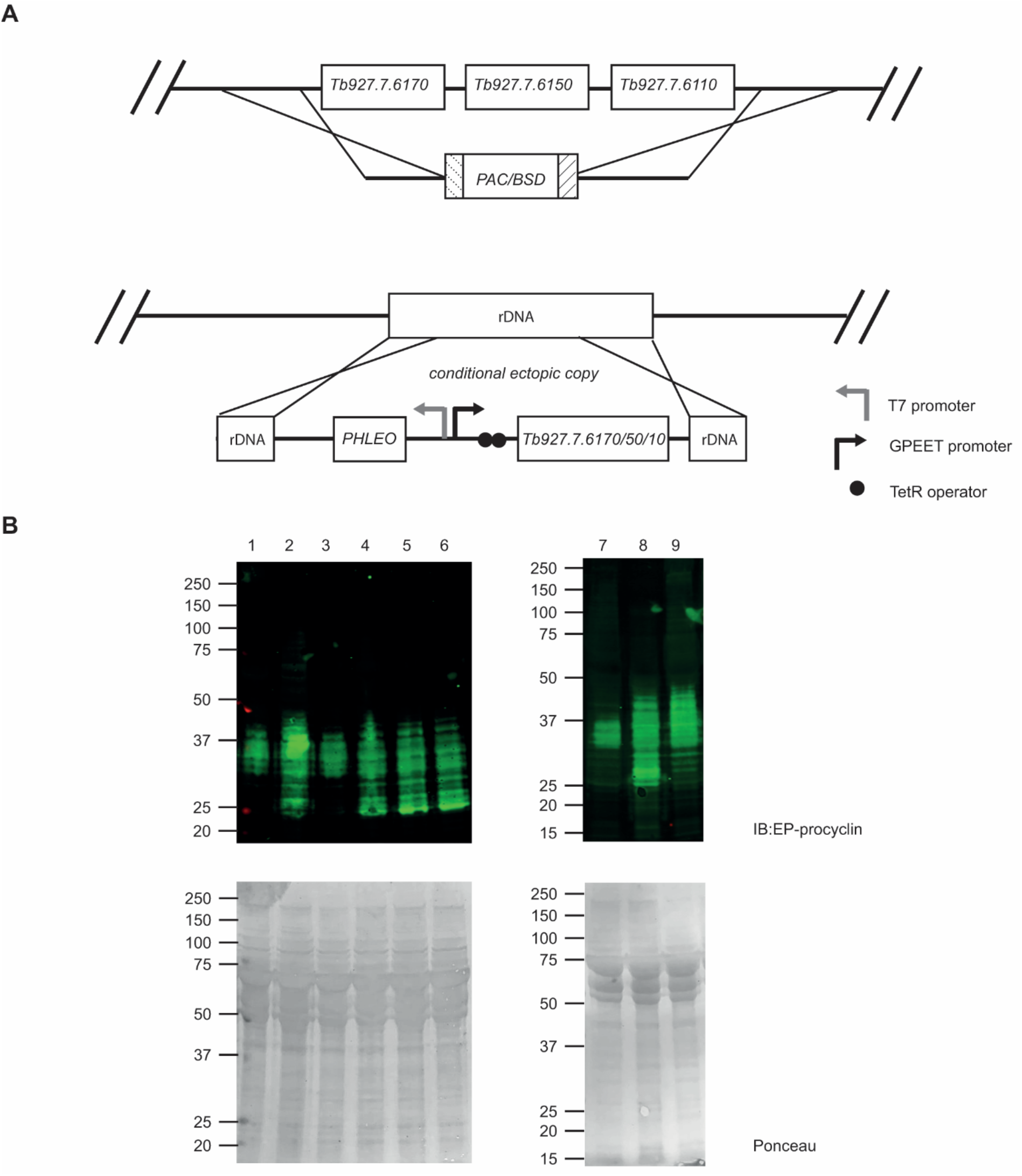
Generation and characterisation of *T. brucei* PCF Tb927.7.6110/6150/6170^-/-^ null and Tb927.7.6110/6150/6170 add back mutants. Schematic of the homologous recombination gene replacement strategy to generate Tb927.7.6110/6150/6170^-/-^ null mutant cells by replacement of the Tb927.7.6110/6150/6170 loci with PAC (puromycin) and BSD (blasticidin) resistance cassettes. Ectopic tetracycline-inducible copies of Tb927.7.6110, 6150 and 6170 (add backs) were introduced into the repetitive ribosomal RNA locus, as indicated. (**B**) Anti-EP procyclin Western blot of samples of PCF WT cells (lanes 1 and 7), Tb927.7.6110/6150/6170^-/-^ null mutant cells (lanes 2 and 8), Tb927.7.6110 add back cells +Tet (lane 3) and -Tet (lane 4), Tb927.7.6150 add back cells +Tet (lane 5) and -Tet (lane 6) and Tb927.7.6170 add back cells +Tet (lane 9). Ponceau staining of blots prior to blocking and antibody staining indicate similar protein loading within each blot.

Using the procedure described in (28) the procyclins purified from wild-type PCF cells and the Tb927.7.6110/6150/6170^-/-^ null mutant were analysed by negative ion MALDI-ToF following aqueous HF dephosphorylation and mild acid treatment. This showed that both the wild-type and null mutant cells were expressing EP-procyclins, predominantly the EP3 form, and not GPEET-procyclins (**Fig. S2**). We therefore probed Western blots of wild-type and mutant PCF cell lysates with anti-EP procyclin antibody. This showed that deletion of the Tb927.7.6110/6150/6170 locus had a significant effect on the mature procyclin profile (**Fig. 3*B***, compare lanes 1 and 2). In the null mutant, the procyclins adopted a more polydisperse appearance ranging from substantially lower to higher apparent MW compared to wild-type procyclins. The deletion of the Tb927.7.6110/6150/6170 locus did not, however, affect cell surface expression of the EP-procyclins as judged by immunofluorescence microscopy (**Fig. S3*A***).

Each of the Tb927.7.6110/6150/6170 genes was individually added back to the Tb927.7.6110/6150/6170^-/-^ null mutant in the form of tetracycline-inducible ectopic copies incorporated into the repetitive ribosomal RNA locus of the parasite using the pLew100 vector (**Fig. 3*A***). The RT-qPCR data for the Tb927.7.6110/6150/6170 transcripts in the wild-type and Tb927.7.6110/6150/6170 add back cells are shown in (**Fig. S3*B***,***C***,***D***).

Activation of an ectopic Tb927.7.6150 gene had no effect on the procyclin SDS-PAGE pattern (**Fig. 3*C***, lane 5) whereas activation of an ectopic Tb927.7.6110 gene reverted the procyclin SDS-PAGE pattern to that of the wild-type (**Fig. 3*C***, lane 3). Activation of an ectopic Tb927.7.6170 gene had an unexpected effect on the procyclin SDS-PAGE pattern (**Fig. 3*B***, lane 9); i.e., the generation of higher apparent MW procyclins (discussed later).

On the basis of these results, we hypothesised that: (i) Tb927.7.6110 alone was sufficient to reverse the Tb927.7.6110/6150/6170^-/-^ null phenotype, and therefore the best candidate for a GPI-PLA2 encoding gene. (ii) Tb927.7.6150 encodes a catalytically inactive variant, consistent with the absence of putative active-site aspartic acid and histidine residues (**Fig. 2*B***). (iii) Tb927.7.6170 might encode a similar activity to Tb927.7.6110, possibly of a GPI-PLA1.

Procyclin preparations from the wild-type, Tb927.7.6110/6150/6170^-/-^ null mutant and the Tb927.7.6110 and Tb927.7.6170 tetracycline-induced add-backs were subjected to nitrous acid deamination and solvent extraction to isolate the released PI components of the procyclin GPI anchors (11) (**Fig. 4*A***). These released PI preparations were analysed by negative ion ES-MS (**Fig. 4*B-E*** and **Table 1**) and by ES-MS^2^ (**Fig. S4**).

**Table 1.** Interpretation of major PI species based on negative ion ES-MS and ES-MS^2^ analysis

| Observed<br>*m/z*<br>[M-H]<sup>-</sup> | Expected<br>*m/z*<br>[M-H]<sup>-</sup> | Molecular<br>formula | *sn*-1 in<br>glycerol<br>backbone | *sn*-2 in<br>glycerol<br>backbone | FA in<br>Inositol ring | WT<br>DM | PLA2<br>KO | 6110<br>+tet | 6170<br>+tet |
| --- | --- | --- | --- | --- | --- | --- | --- | --- | --- |
| 837.55062* | 837.54985 | C43H83O13P1 | C18:0-acyl | - | C16:0 | + | + | + | + |
| 847.57119 | 847.57058 | C45H85O12P1 | C18:0-alkyl | - | C18:2 | + | + | + | + |
| 861.55062* | 861.54985 | C45H83O13P1 | C18:0-acyl | - | C18:2 | + | + | + | + |
| 863.56457 | 863.56550 | C45H85O13P1 | C18:0-acyl | - | C18:1 | + | + | + | + |
| 865.58040 | 865.58115 | C45H87O13P1 | C18:0-acyl | - | C18:0 | + | + | + | + |
| 1073.76609 | 1073.76386 | C59H111O14P | C18:0-acyl | C18:1-acyl | C14:0 |  | + |  | + |
| 1085.80368 | 1085.80025 | C61H115O13P | C18:0-alkyl | C18:2-acyl | C16:0 |  | + |  | + |
| 1099.78246 | 1099.77951 | C61H113O14P | C18:0-acyl | C18:2-acyl | C16:0 |  | + |  | + |
| 1101.79769* | 1101.79516 | C61H115O14P | C18:0-acyl | C18:1-acyl | C16:0 |  | + |  | + |
| 1125.79727* | 1125.79516 | C63H115O14P | C18:0-acyl | C18:1-acyl | C18:2 |  | + |  | + |
| 1141.79110 | 1141.82646 | C64H119O14P | C19:0-acyl | C18:0-acyl | C18:2 |  | + |  | + |
| 1157.78572 | 1159.87341 | C65H125O14P | C20:0-acyl | C18:1-acyl | C18:0 |  | + |  | + |

+ indicates the presence of respective *m/z* ion the sample.
\*Refer to Supplementary **Fig. S2** for the corresponding ES-MS<sup>2</sup>

**Figure 4:**
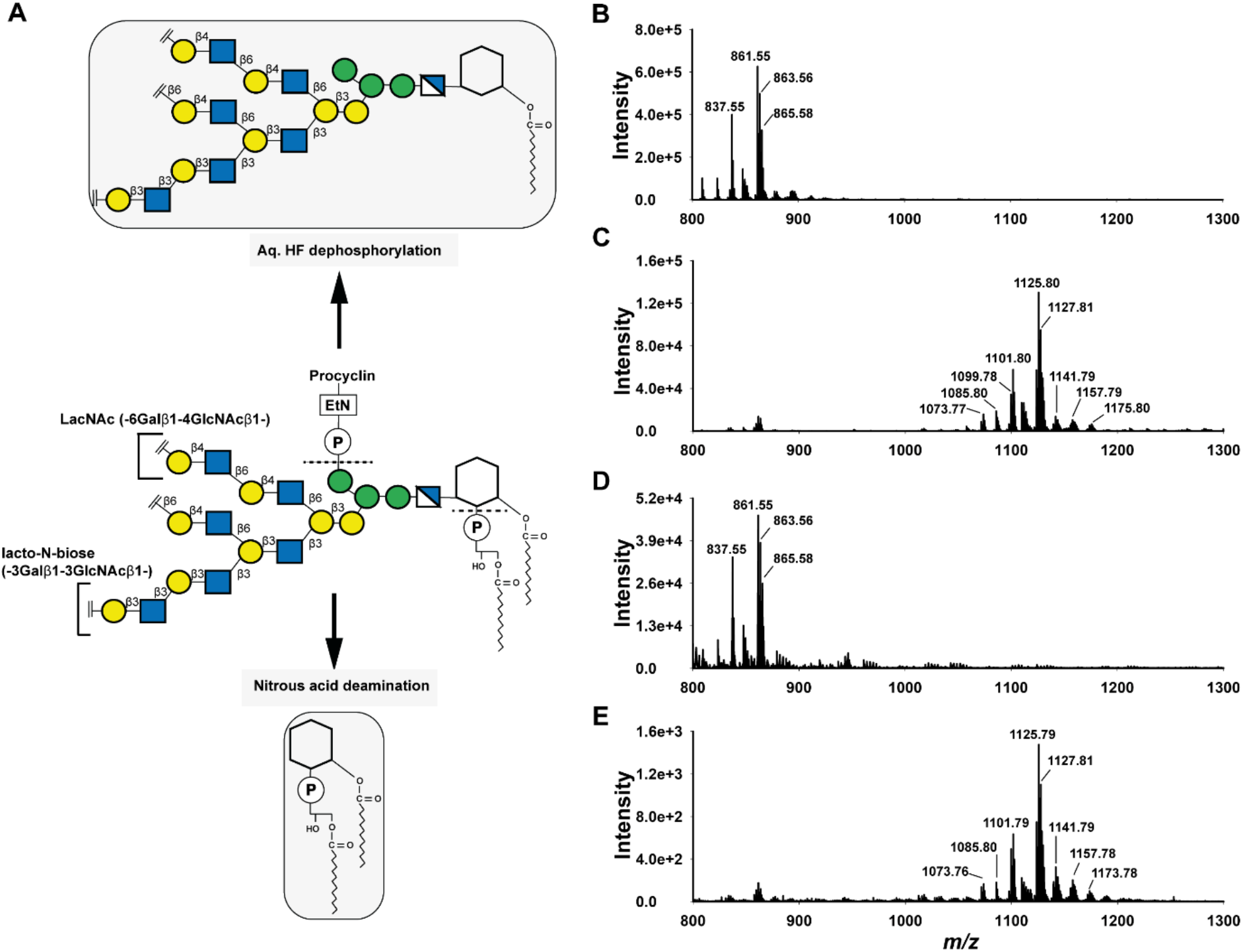
Analysis of procyclin PI species. (**A**) Schematic of the GPI anchor structure core of procyclin (centre) showing the Lacto-N-biose (−3Galβ1-3GlcNAc β1-) and LacNAc (−6Galβ1-4GlcNAc β1-) repeats in the GPI glycan side chains. The schematic also shows the expected liberation of PI structures from GPI anchored procyclin after nitrous acid deamination (bottom) and the expected GPI glycan core released after aq. HF dephosphorylation (top). (**B – E)** show the negative ion ES-MS spectra of the PI species released from wild-type (**B**), Tb927.7.6110/6150/6170^-/-^ null mutant (**C**), Tb927.7.6110^+^ add back (**D**) and Tb927.7.6170^+^ add back (**E**) procyclins. The PI species were observed as [M-H]^-^ ions and the major [M-H]^-^ precursor ions were subjected to ES-MS^2^ using collision induced dissociation (Supplementary **Fig. S2**). Refer to **Table 1** for the fatty acid compositions of various PI ions.

The compositions (by accurate mass) and structures (deduced from the MS^2^ product ion spectra) of the wild-type PIs were as expected from previous reports (12). Thus, the wild-type PI structures were all *lyso*-PI structures with a mixture of fatty acids ester-linked to the inositol ring (**Fig. 4*B*, Fig. S4A, Table 1**). In contrast, the major released PI structures from the Tb927.7.6110/6150/6170^-/-^ null mutant were mostly diacyl-PIs with the same range of fatty acids attached to the inositol ring (**Fig. 4*C*, Fig. S4*B*, Table 1**). These data show unambiguously that one or more genes in the Tb927.7.6110/6150/6170 locus is/are required for GPI-PLA2 activity in PCF cells.

The add back of Tb927.7.6110 to the Tb927.7.6110/6150/6170^-/-^ null mutant also reverted the pattern of procyclin released PIs to that of the wild-type (**Fig. 4*D***), demonstrating that Tb927.7.6110 alone is sufficient for GPI-PLA2 activity in PCF cells. The add back of Tb927.7.6170 had no effect on the Tb927.7.6110/6150/6170^-/-^ null mutant procyclin PI structures (**Fig. 4*E***).

From these data we conclude that the Tb927.7.6110 gene alone is necessary and sufficient for TbGPI-PLA2 activity in PCF cells. Further, due to its structural similarity with mammalian PGAP6 we postulate that it directly encodes TbGPI-PLA2 activity.

### Effects of Tb927.7.6110/6150/6170 ablation and Tb927.7.6110 and Tb927.7.6170 re-expression on procyclin GPI glycan structure

The procyclin preparations used to analyse the nitrous acid released PI species (**Fig. 4, Table 1**) were also subjected to neutral monosaccharide composition analysis by GC-MS and, following dephosphorylation with aqueous HF and permethylation, to MALDI-ToF and methylation linkage analysis by GC-MS.

In the monosaccharide composition analysis, a reduction in Gal and GlcNAc relative to Man was observed in the procyclins from the Tb927.7.6110/6150/6170^-/-^ null mutant compared to wild-type procyclins (**Fig. 5*A*** and **5*B***). This reduction in Gal : Man ratio was largely recovered in the Tb927.7.6110 and Tb927.7.6170 add back procyclins (**Fig. 5*C,D***).

**Figure 5:**
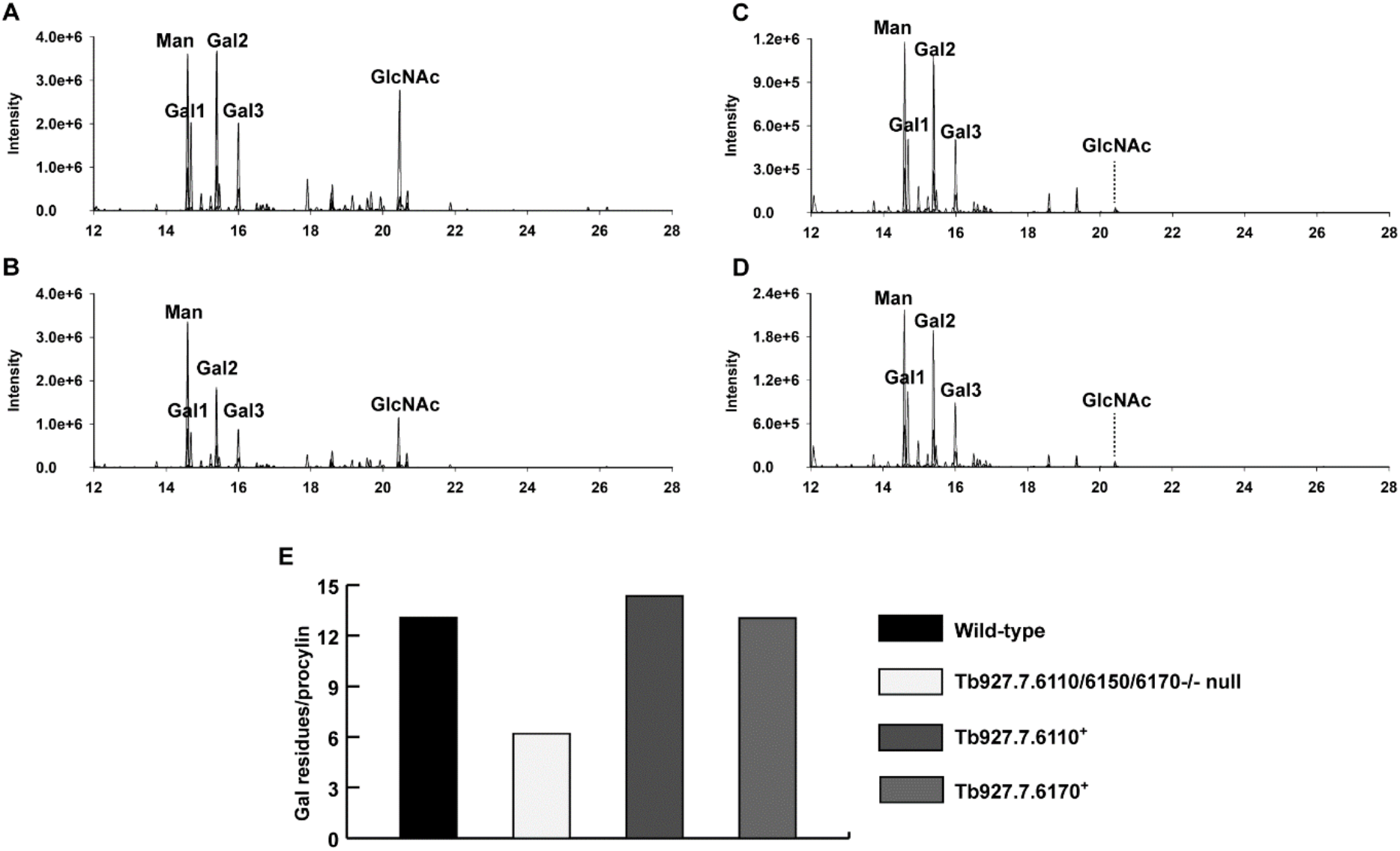
Monosaccharide analyses of wild-type and mutant procyclins. Extracted ion GC-MS chromatograms of wild-type (**A**), Tb927.7.6110/6150/6170^-/-^ null mutant (**B**), Tb927.7.6110^+^ add back (**C**) and Tb927.7.6170^+^ add back (**D**) carbohydrate analyses. (**E**) Average Gal content per procyclin molecule for wild-type, Tb927.7.6110/6150/6170^-/-^ null mutant, Tb927.7.6110^+^ add back and Tb927.7.6170^+^ add back procyclins.

We can use the the Gal : Man ratio as an indicator of how large, on average, the GPI anchor glycan sidechains are. Each EP-procyclin molecule contains a Man_5_GlcNAc_2_ N-linked glycan and an ethanolamine-P-6Man*α*1-2Man*α*1-6Man*α*1-4GlcN GPI glycan core (11). Of the latter, only two of the three Man residues are detected by the GC-MS method employed (because of the acid-stability of the 6-linked phosphate group) (2). This means that there are 7 detectable Man residues per molecule of procyclin. Based on this, we estimate the average number of Gal residues in the GPI anchor sidechains of the wild-type and add-back procyclins as 12-14 and in the Tb927.7.6110/6150/6170^-/-^ null mutant as 6-7 (**Fig. 5*E***).

We focussed on comparing the GPI anchor glycan sidechains in the wild-type and Tb927.7.6110/6150/6170^-/-^ null mutant procyclins. The positive ion MALDI-ToF spectra of the aq. HF released and permethylated GPI glycans were consistent with a smaller average size of GPI glycan sidechain in the mutant procyclins (**Fig. 6**).

**Figure 6:**
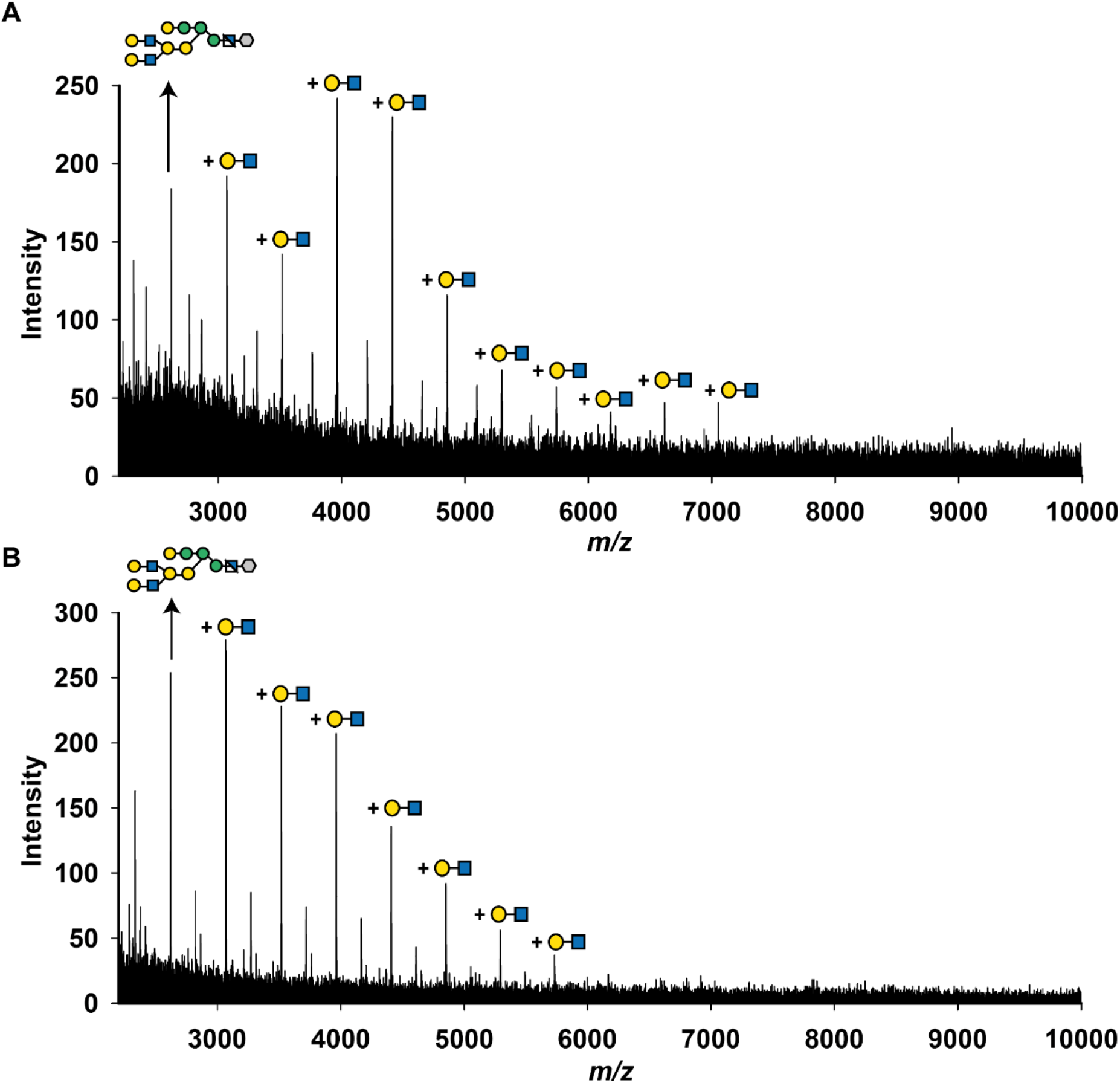
GPI glycan analysis after permethylation: The GPI glycans of wild-type and Tb927.7.6110/6150/6170-/-null mutant procyclins were released by aq. HF treatment (see Fig. 4A) and subjected to permethylation, which simultaneously removes the inositol-linked fatty acid. The permethylated GPI glycans of wild-type (**A**) and Tb927.7.6110/6150/6170-/-null mutant (**B**) were analysed by positive ion MALDI-ToF. A series of GPI-glycans with the addition of Hex-HexNAc repeats (449 Da difference, equivalent to permethylated Hex-HexNAc) are indicated.

The GC-MS methylation linkage analysis results for these same preparations were normalised to the areas for the non-reducing terminal-Man residues arising from the common Man_5_GlcNAc_2_ N-linked glycans. As expected, the relative 3,6-disubstituted Man signals, also arising from the common Man_5_GlcNAc_2_ N-linked glycans, were very similar. On the other hand, the relative amounts of Gal and GlcNAc derivatives were significantly reduced for the Tb927.7.6110/6150/6170^-/-^ null mutant sample, particularly those corresponding to terminal-Gal, 3-substituted Gal, 3,6-disubstituted Gal and 3-substituted GlcNAc residues (**Fig. 7, Table S1**). These data suggest that the Tb927.7.6110/6150/6170^-/-^ null mutant GPI glycan sidechains are principally deficient in lacto-N-biose (−3Gal*β*1-3GlcNAc*β*1-) repeats.

**Figure 7:**
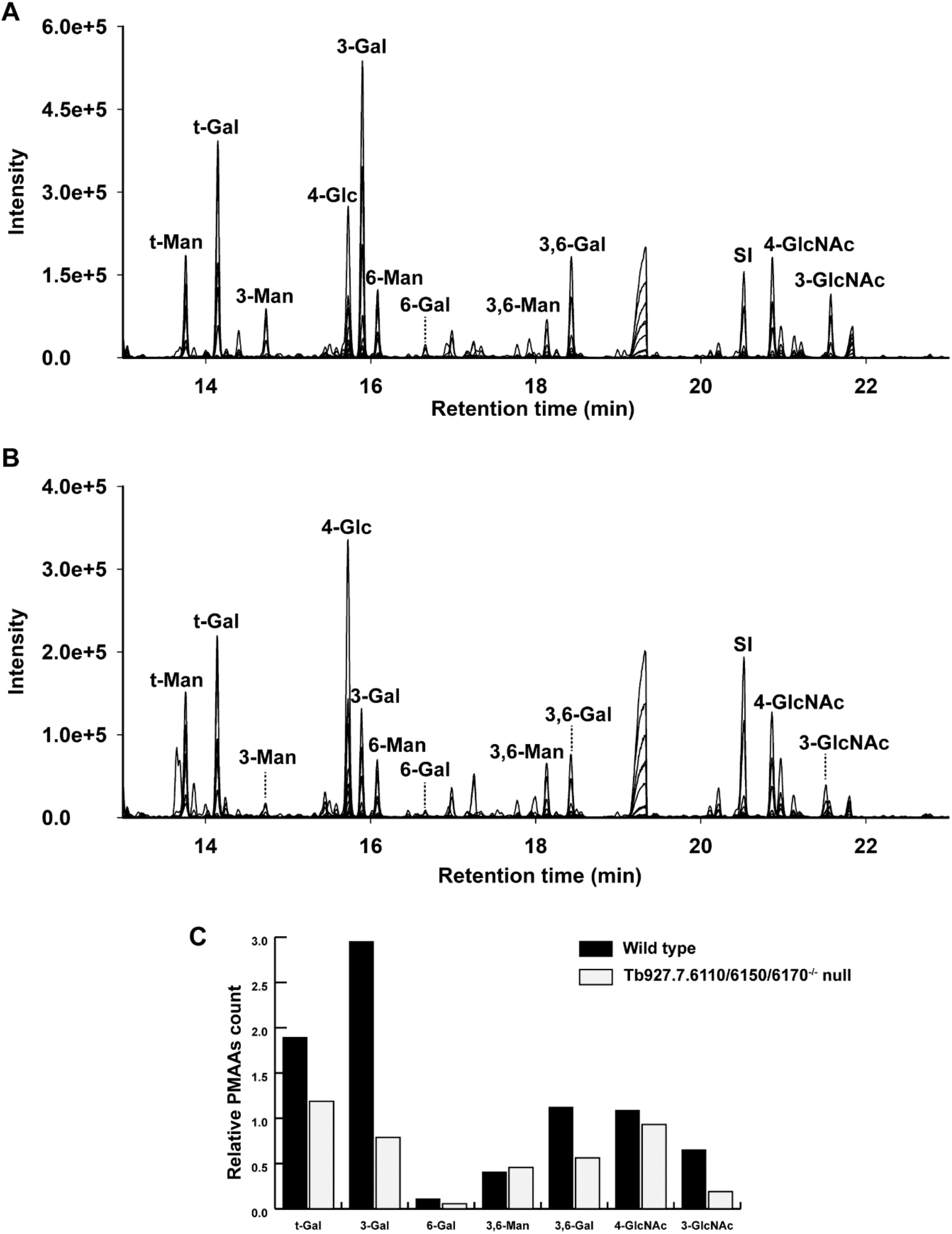
Methylation linkage analysis of GPI glycans. The permethylated GPI glycans of wild-type (**A**) and Tb927.7.6110/6150/6170-/-null mutant samples were hydrolysed, reduced, acetylated and the resulting PMAA derivatives were analysed by GC-MS. The peak marked as SI is an internal standard of *scyllo*-inositol hexa-acetate. The chromatograms shown above are merged extracted ion chromatograms for characteristic PMAA fragment ions (*m/z* 102, 117, 118, 129, 145, 159, 161, 162, 168, 189, 190, 205, 210, 233, and 234). The PMAA peaks are annotated according to the original substitution pattern of the monosaccharides in the native glycans. For example, t-Man refers to non-reducing-terminal mannose and 3-Gal refers to 3-O-substituted galactose (see **Table S1**). (**C**) Represents the comparison of PMAA signals of wild-type and Tb927.7.6110/6150/6170^-/-^ null mutant samples, normalised to non-reducing-terminal mannose (t-Man) in each sample.

## Discussion

The identity of the GPI-PLA2 enzyme responsible for GPI precursor fatty acid remodelling in *T. brucei* (**Fig. 1**) has been elusive for many years, but mammalian PGAP6 (24, 25) provided an inroad to identify potential GPI-PLA2 encoding genes in *T. brucei*. Three PGAP6-related genes were identified (**Fig. 2**) in tandem array with homologous intergenic regions, making individual gene knockouts difficult to achieve. We deleted all three in PCF *T. brucei* to make a Tb927.7.6110/6150/6170^-/-^ null mutant and observed a biochemical phenotype consistent with the deletion of GPI-PLA2 activity: Namely, the presence of *sn*-1,2-diacylglycerol in place of *sn*-1-acyl-2-*lyso*-glycerol in the PI component of the GPI membrane anchors of the procyclin surface glycoproteins. Through a tetracycline-inducible add-back approach, we were able to show that only one of the three genes, Tb927.7.6110, was necessary and sufficient to reverse the mutant phenotype to that of wild-type. From this, we conclude that Tb927.7.6110 encodes the GPI-PLA2 of GPI precursor fatty acid remodelling, a biochemical activity originally described by Masterson *et al* in (16).

We had expected that deletion of GPI-PLA2 activity might result in slightly increased mobility of the procyclins on SDS-PAGE because a more hydrophobic (diacyl-PI containing) GPI anchor lipid should bind more SDS than the native (acyl-*lyso*-PI containing) GPI lipid. However, we were surprised at the observed increase in mobility and increase in dispersion (**Fig. 3**). Since the apparent MW heterogeneity of procyclins stems primarily from the heterogeneity of their GPI anchor carbohydrate sidechains (12, 28), we compared the carbohydrate contents and

GPI glycan structures of the wild-type and Tb927.7.6110/6150/6170^-/-^ null mutant PCF cells. The data showed an approximate halving in Gal content in the mutant (**Fig. 5**) and a reduction in the size profile of GPI glycans released from the procyclins (**Fig. 6**), apparently from the selective loss of lacto-N-biose (−3Gal*β*1-3Gal*β*1-) repeat units (**Fig. 7**). We therefore interpret the change in SDS-PAGE pattern of the procyclins in the Tb927.7.6110/6150/6170^-/-^ null mutant as a combination of a reduction in SDS-binding capacity and in overall GPI glycan sidechain size.

The change in overall GPI sidechain size in the Tb927.7.6110/6150/6170^-/-^ null mutant is interesting. One explanation could be that, with an extra fatty acid now present on the GPI anchor, the procyclins and the GPI anchor core present different aspect to the membrane. This might, in turn, reduce accessibility GPI anchor to some of the GPI sidechain processing glycosyltransferases, some or all of which are in the Golgi apparatus, reviewed in (29), reducing overall glycosylation. Another, not mutually exclusive, explanation could be that, with the additional fatty acid present, the transit time of the procyclins through the Golgi might be reduced, leading to less processing. Yet another possibility is that Tb927.7.6110 not only encodes GPI-PLA2 activity but also couples newly synthesised procyclins to their GPI processing glycosyltransferases in some way. This might also explain the odd phenotype of the Tb927.7.6170 add-back where there is no restoration of GPI-PLA2 activity (**Fig. 4**) yet its overexpression causes a notable change in procyclin apparent MW range by SDS-PAGE (**Fig. 3**). It is conceivable, therefore, that the GPI side chain modification is influenced in some way by the interaction of Tb927.7.6110 and Tb927.7.6170 with downstream GPI processing enzymes. Interestingly, a similar inference to this was made recently in BSF *T. brucei* for TbGPI2, a component of the UDP-GlcNAc : PI *α*1-6 GlcNAc transferase complex that catalyses the first committed step in GPI biosynthesis in *T. brucei* (30).

While Tb927.7.6170 does not encode a GPI precursor fatty acid remodelling GPI-PLA2 activity, it might encode a similar activity, such as GPI-PLA1. The latter does not act in PCF cells but is required to complete fatty acid remodelling in BSF cells (**Fig. 1**). The obligate requirement for GPI-PLA2 action and Gup1-mediated myristoylation of the *sn*-2-position prior to GPI-PLA1 action in the BSF GPI fatty acid remodelling pathway (11, 16) make this difficult to assess. Indeed, our attempts to make comparable Tb927.7.6110/6150/6170^-/-^ null and conditional add-back cell lines in BSF *T. brucei* have been unsuccessful.

Another potential role for Tb927.7.6170 could be as a distinct GPI-PLA2 for the process of BSF GPI anchor fatty acid proof-reading, as described in (31, 32). In this process, it is postulated that the myristic acids of the *sn*-1,2-dimyristoylglycerol component of the VSG GPI anchor are removed and replaced by fresh myristic acid in the endosomal compartment of the cell, requiring GPI-PLA2 and GPI-PLA1 activities.

The enzymes of GPI fatty acid remodelling were thought to be potential drug targets for human and animal African trypanosomiases (33). However, these studies with myristate analogues may, in retrospect, have been confounded by the essentiality of trypanosome protein N-myristoyltransferase (NMT) (34–36). More work is required to assess the essentiality of GPI-PLA2 (Tb927.7.6110) in BSF *T. brucei*, and the role and essentiality of its relative Tb927.7.6170 and of their presumably enzymatically inactive relative Tb927.7.6150.

## Experimental procedures

### Cultivation of Trypanosomes

*Trypanosoma brucei brucei* Lister strain 427 procyclic form (PCF) parasites, that maintain T7 polymerase and tetracycline repressor protein under G418 and hygromycin antibiotic selection, were used in this study (37). This genetic background will be referred to from hereon as wild-type. The cells were cultivated in SDM-79 medium supplemented with 15% fetal bovine serum (FCS), Glutamax and hemin, and containing 15 μg/ml G418 and 50 μg/ml hygromycin at 28 °C in a 5% CO_2_ incubator.

### DNA Isolation and Manipulation

Plasmid DNA was purified from *Escherichia coli* DH5α competent cells using a Qiagen Miniprep kit. Extraction and purification of DNA from gels was performed using Qiaquick kits (Qiagen). Custom oligonucleotides were obtained from Thermo Fisher. *T. brucei* genomic DNA was isolated from ∼2 × 10^8^ bloodstream form cells using a DNeasy Blood & Tissue Kit (Qiagen).

### Generation of gene replacement constructs

A full list and descriptions of primers used in this study are summarised in (Supplementary **Table S2)**. A construct containing a puromycin acetyltransferase (*PAC*) gene flanked at the 5’ end with 500 bp of Tb927.7.6170 5’ UTR following by 135 bp of *T. brucei* actin gene 5’ UTR, and at the 3’ end with 293 bp actin 3’ UTR followed by 500 bp of Tb927.7.6110 3’ UTR was synthesized by Genscript. The same construct containing a blasticidin-S deaminase (*BSD*) gene in place of *PAC* was generated by Gibson assembly (NEB) using primers ZJ1-ZJ4 (**Table S2**). The first copy of the Tb927.7.6170, Tb927.7.6150 and Tb927.7.6110 locus was replaced with the *PAC* drug resistance construct to generate a single knock out (sKO) mutant for these three genes. The second copy of the Tb927.7.6170, Tb927.7.6150 and Tb927.7.6110 locus was replaced by the *BSD* resistance construct to generate the Tb927.7.6170, Tb927.7.6150, Tb927.7.6110^-/-^ double knockout (dKO) mutant. The identity of all constructs was confirmed by DNA sequencing.

### Generation of *T. brucei* add-back constructs

To generate the add back constructs for three genes (Tb927.7.6110, Tb927.7.6150, Tb927.7.6170), their individual open reading frames (ORFs) were synthesized by Genscript. The individual ORFs were then amplified by PCR using primers ZJ6-ZJ10 (**Table S2**) and cloned individually into a pLEW100v5 vector (pLEW100v5 was a gift from George Cross; Addgene plasmid # 24011) using Gibson assembly, placing the genes under a tetracycline inducible promoter. The identity of all constructs was confirmed by DNA sequencing.

### Transformation of *T. brucei* procyclic form cells

The gene replacement constructs, and ectopic expression (add back) constructs were purified using the Qiagen Miniprep kit, linearised by NotI (NEB) restriction digestion, precipitated and washed twice with 70% ethanol and re-dissolved in sterile water. The precipitated linearised DNA was used to electroporate *T. brucei* PCF as described in (37). The generation of sKO and dKO (null) mutants was confirmed by Southern blot.

### Southern blotting

The DNA probes used in this study were digoxigenin (DIG)-labelled. These were generated using a PCR DIG Probe Synthesis Kit (Roche) according to the manufacture’s protocol. ZJ13 and ZJ14 primers were used for synthesis of the *PAC* resistance cassette probe, ZJ15 and ZJ16 primers were used for synthesis of the *BSD* resistance cassette probe, ZJ17 and ZJ18 primers were used for synthesis of the Tb927.7.6170 ORF probe. For analysis of *T. brucei* mutants, 5 µg of genomic DNA from the clones were digested solely with XhoI (NEB) overnight at 37 °C. RNAase (Sigma) was added to the reaction to avoid RNA contamination for later detection. Endonuclease digested gDNA was separated on a 0.8% (w/v) agarose gel for 4 h at 40 V in TAE buffer. The gel was washed with 0.25 M HCl with mild agitation (30 rpm) for 10 min to depurinate the DNA following by 15 min denaturation with 0.5 M NaOH and 20 min neutralisation with buffer 1 M Tris-HCl, 1.5 M NaCl, pH 7.5. After these steps, the DNA samples were transferred to a positively charged nylon membrane (Roche) through reverse capillary action overnight with 10 × saline-sodium citrate buffer (SSC) buffer. Transferred DNA fragments were covalently cross-linked to the membrane by UV cross-linking in a CL-100 (UVP) UV crosslinker at 1200 mJoules. The membrane was pre-incubated with 20 mL DIG Easy Hyb™ Granules solution (Roche) at 42 °C for 1 h in a hybridisation oven (Techne™ Hybridisation Oven). The DIG-labelled probe was denatured for 5 min at 100 °C following by a rapid cool down on ice prior to mixing with 20 mL DIG Easy Hyb™ Granules solution. The membrane was incubated overnight with this hybridisation solution with 20 µL denatured probe. Following the hybridisation, the membrane was washed twice at low-stringency (42 °C, 5 min, 1 × SSC with 0.01% (w/v) SDS) and high-stringency (65 °C, 15 min, 0.5 × SSC with 0.01% (w/v) SDS) sequentially. The blot was then developed with DIG wash and block buffer set (Roche) following the manufacturer’s instructions. The blots were equilibrated in 1 × wash buffer for 5 min at room temperature before being blocked in blocking buffer for 30 min. The anti-DIG AP-conjugate antibody (Roche) was then added to the blocking buffer with a 1:10,000 dilution and the blot was incubated further for 30 min followed by twice washing with 1 × wash buffer (15 min). The membrane was placed in a plastic folder and chemiluminescent substrate CSPD (Roche) was applied and incubated for the detection. The blots were exposed onto Amersham Hyperfilm™ ECL film for 1-30 min and developed with a KODAK film developer. For stripping the blot, 0.4 M NaOH was applied twice to the membrane and for 5 min at 42 °C in the hybridisation oven. The blot was then washed three times with 1 x SSC buffer for 10 min at 42 °C before re-probing.

### Quantitative Real-Time PCR (qRT-PCR)

RNA was extracted from 1 × 10^7^ *T. brucei* PCF wild-type, *Tb927*.*7*.*6110/6150/6170*^-/-^ null mutant and cells overexpressing *Tb927*.*7*.*6170, 6150* and *6110* using RNeasy Plus MiniKit (Qiagen). RT-qPCR was performed using the Luna Universal qPCR Master Mix (NEB) on a QuantStudio 3 Real-Time PCR System (Applied Biosystems) according to manufacturer’s instructions. Primers (ZJ19-24) used for amplifying the specific regions of *Tb927*.*7*.*6170/*.*6150/ 6110* are shown in (**Table S2**). The individual specific gene transcript levels are presented as fold change relative to wild type controls, determined using the comparative CT (ΔΔCT) method using β-tubulin as an endogenous control for normalisation. All reactions were carried out in technical triplicates.

### Western blotting of *T. brucei* whole cell lysate

For Western blot analysis, 5 × 10^6^ – 1 × 10^7^ cells were lysed and solubilised in 1x SDS sample buffer containing 0.1 M DTT by heating at 55 °C for 20 mins. The proteins were resolved by SDS-PAGE (approximately 1 × 10^7^ cell equivalents/lane) on NuPAGE bis-Tris 4–12% gradient acrylamide gels (Invitrogen) and transferred to nitrocellulose membrane (Invitrogen). Ponceau S staining was used as transfer control and to confirm the equal loading. Procyclins were detected using a monoclonal anti-EP antibody (1:750 dilution) in blocking buffer (50 mM Tris-HCl pH 7.4, 0.15 M NaCl, 0.25% BSA, 0.05% (w/v) Tween-20, 0.05% NaN3 and 2% (w/v) Fish Skin Gelatin). Detection was performed IRDye® 800CW Goat anti-Mouse at 1:15,000 in blocking buffer. The immunoblot was analysed on the LI-COR Odyssey Infrared Imaging System (LICOR Biosciences).

### Extraction and purification of procyclins

Procyclins were purified from 10^10^ cells by organic solvent extraction and octyl-Sepharose chromatography as previously described (30). Briefly, the cells were extracted three times with chloroform/methanol/water (10:10:3, v/v). The pellet obtained after the delipidation process was extracted twice with 9% butan-1-ol in water. The pooled supernatants were back-extracted twice with an equal volume of 9% water in butan-1-ol and the lower 9% butan-1-ol in water phase containing procyclins was recovered and dried under N_2_ stream.

Solvent extracted procyclins were used for nitrous acid deamination and monosaccharide composition analysis. Whereas, for analysing the procyclin by MALDI-ToF and permethylated GPI glycans, the extracted procyclins were further purified using octyl-Sepharose 4B (Sigma) chromatography. Briefly, the extracted procyclins, dried and redissolved in buffer A (5% propan-1-ol in 0.1 M ammonium acetate) were applied to 0.5 ml of octyl-Sepharose 4B, packed in a disposable column and pre-equilibrated with buffer A. The column was washed with buffer A followed by buffer B (5% propan-1-ol). The procyclins were then eluted in buffer C (50% propan-1-ol), concentrated and dried until further use.

### Analysis of procyclins by MALDI-ToF

The solvent extracted procyclins (approx. 600 pmole) were dried and subjected to dephosphorylation using 50 μl of ice-cold 50 % aqueous hydrogen fluoride (aq. HF) for 24 h at 0 °C to cleave the GPI anchor ethanolamine-phosphate bond. Dephosphorylated samples were further subjected to mild acid treatment with 50 μl of 40 mM trifluoroacetic acid at 100 °C for 20 min to cleave Asp-Pro bonds and thus remove the N-glycosylated N-termini. The samples were dried and redissolved in 5 μl of 0.1% trifluoroacetic acid and the aliquots were co-crystallised with α-cyano-4-hydroxycinnamic acid matrix (10 mg/ml in 50% acetonitrile, 0.1% trifluoroacetic). The samples were analysed by linear-mode negative-ion MALDI-ToF (Autoflex Speed MALDI-ToF MS system by Bruker).

### Deamination of extracted procyclin and ES-MS analysis

The procyclins were deaminated as described (12) with minor modification. Briefly, 15 % of the extracted procyclin preparation was dried and deaminated with 50 µl of 0.3 mM sodium acetate buffer (pH 4.0) containing 250 mM sodium nitrite for 1 h at room temperature. The samples were replenished with a further 50 µl of 0.3 mM sodium acetate buffer (pH 4.0) containing 250 mM sodium nitrite and incubated for another 2 h at 37 °C. The deaminated procyclins were extracted three times with water saturated butan-1-ol (i.e., butan-1-ol containing 9 % water). The upper butan-1-ol phase containing the released phosphatidylinositol (PI) moieties were pooled in a fresh tube and dried under N_2_. To analyse the released PI components, the samples were redissolved in 100 µl of chloroform/ methanol (1:1, v/v) and infused into an LTQ Orbitrap Velos Pro mass spectrometer (Thermo Scientific) using static infusion nanoflow probe tips (M956232AD1-S, Waters). Data were collected in negative ion mode for ES-MS and ES-MS^2^. Negative ion spray voltage was 0.8 kV, the temperature of ion transfer tube was 275 °C and collision induced dissociation (CID) was used for MS^2^ fragmentation, using 25-35% collision energy.

### GC-MS Monosaccharide composition

Aliquots (3%) of the extracted procyclins were mixed with 1 nmol *scyllo*-inositol internal standard and subjected to methanolysis, re-N-acetylation and trimethylsilylation (38). The resulting 1-O-methyl-glycoside TMS derivatives were analysed by GC-MS (Agilent Technologies, 7890B Gas Chromatography system with 5977A MSD, equipped with Agilent HP-5ms GC Column, 30 m X 0.25 mm, 0.25 µm). To analyse the data, the total ion chromatogram (TIC) was extracted for characteristic ions (*m/z* 204, *m/z* 217, *m/z* 173, *m/z* 305, *m/z* 318).

### Permethylation and analysis of GPI glycans by MALDI-ToF MS

The octyl-Sepharose purified procyclin samples were subjected to dephosphorylation and permethylation as described in (30). Briefly, the procyclins were treated with 100 µl of ice-cold 50% aqueous hydrogen fluoride (aq. HF) for 24 h at 0 °C to cleave the GPI anchor ethanolamine-phosphate-mannose and inositol-phosphate-acylglycerol phosphodiester bonds and release the GPI glycan. After removing the aq. HF by freeze drying, the released GPI glycans were resuspended in 100 µl water, centrifuged at 16000 x g for 10 min and obtained in the supernatant. The permethylation of released GPI glycans was performed using the sodium hydroxide method, as described earlier (38) to obtain the GPI glycans bearing a fixed positive charge in the form of an N-trimethyl-glucosamine quaternary ammonium ion. Aliquots (10%) of permethylated GPI-glycan were analysed using positive ion MALDI-ToF MS (Autoflex Speed MALDI-ToF MS system by Bruker). The permethylated GPI glycans samples were co-crystallised with a matrix of 2,5-dihydroxybenzoic acid (20 mg/ ml 30 % acetonitrile and 0.1 % TFA) and analysed in reflectron positive ion mode.

### Methylation linkage analysis

Methylation linkage analysis of GPI glycans was performed as described (30). Briefly, 80 % of the permethylated GPI glycan samples were subjected to acid hydrolysis, NaB[^2^H]_4_ reduction, and acetylation to generate partially methylated alditol acetates (PMAAs) which were analysed by GC-MS.

## Abbreviations

The abbreviations used are

BSF: bloodstream form
PCF: procyclic form
GPI: glycosylphosphatidylinositol
HYG: hygromycin phosphotransferase
PAC: puromycin acetyltransferase
BSD: blasticidin S deaminase
LacNAc: N-acetyllactosamine
VSG: variant surface glycoprotein
ER: endoplasmic reticulum
GlcN: glucosamine
Man: mannose
Gal: galactose
GlcNAc: N-acetylglucosamine
PI: phosphatidylinositol
ES-MS: electrospray-mass spectrometry
MALDI-ToF: matrix assisted laser desorption ionization-time of flight
GC-MS: gas chromatography-mass spectrometry
PMAAs: partially methylated alditol acetates.

## Acknowledgments

This work was supported by a China Scholarship Council PhD scholarship to Z. J. (201706310166) and a Wellcome Investigator Award to M.A.J.F. (101842/Z13/Z).

## CRediT Author Statement

**Zhe Ji:** Conceptualization, Data Curation, Formal Analysis, Investigation, Methodology, Writing - Original Draft. **Rupa Nagar:** Data Curation, Formal Analysis, Investigation, Methodology, Writing - Original Draft. **Samuel M. Duncan:** Supervision, Resources. **Maria Lucia Sampaio Guther:** Supervision, Resources. **Michael A.J. Ferguson:** Conceptualization, Data Curation, Formal Analysis, Funding Acquisition, Methodology, Writing - Original Draft, Writing – Review and Editing.

## Notes

The authors declare that they have no conflicts of interest with the contents of this article.

### Competing Interest Statement

The authors have declared no competing interest.

