## Supplemental Data for "Identification of the glycosylphosphatidylinositol-specific phospholipase A2 (GPI-PLA2) of GPI fatty acid remodelling in *Trypanosoma brucei*"

### Supporting information

**Table: S1**: GC-MS methylation linkage analysis of GPI-glycans from wild-type and Tb927.7.6110/6150/6170$^{-/-}$ null mutant cells

**Table S2**: List of oligonucleotides primers used in this study

**Figure S1**: Verification of the Tb927.7.6110/6150/6170$^{-/-}$ null mutant genotype by Southern blot.

**Figure S2**: Identification of procyclin types by MALDI-ToF

**Figure S3**: Immunofluorescence microscopy for surface localisation of procyclins in PCF wild-type and Tb927.7.6110/6150/6170$^{-/-}$ null mutants and RT-qPCR for ectopic copy expression.

**Figure S4**: ES-MS$^2$ identification of PI species

**Table S1: GC-MS methylation linkage analysis of GPI-glycans from wild-type (WT) and Tb927.7.6110/6150/6170$^{-/-}$ null (Null$^{-/-}$) mutant cells.** The GPI glycans were permethylated, hydrolysed, deutero-reduced, and acetylated to yield PMAAs for analysis by GC-MS. Residue types were deduced from the electron-impact mass spectra and retention times.

| **PMAA derivative** | **Residue types** | **RT [a] (min)** | **EIC ion used[b]** | **Sample[c]** | |
|---|---|---|---|---|---|
| | | | | **WT** | **Null$^{-/-}$** |
| [1-$^2$H]-1,5-Di-*O*-acetyl-2,3,4,6-tetra-*O*-methylmannitol | t-Man | 13.75 | 102 | 1.0 | 1.0 |
| [1-$^2$H]-1,3,5-Tri-*O*-acetyl-2,4,6-tri-*O*-methylmannitol | 3-Man | 14.73 | 118 | 0.5 | 0.7 |
| [1-$^2$H]-1,5,6-Tri-*O*-acetyl-2,3,4-tri-*O*-methylmannitol | 6-Man | 16.08 | 102 | 0.67 | 0.45 |
| [1-$^2$H]-1,3,5,6-Tetra-*O*-acetyl-2,4-di-*O*-methylmannitol | 3,6-Man | 18.13 | 118 | 0.41 | 0.45 |
| [1-2H]-1,5-Di-*O*-acetyl-2,3,4,6-tetra-*O*-methylgalactitol | t-Gal | 14.14 | 102 | 1.89 | 1.18 (37 % down) |
| [1-2H]-1,3,5-Tri-*O*-acetyl-2,4,6-tri-*O*-methylgalactitol | 3-Gal | 15.89 | 118 | 2.95 | 0.78 (73 % down) |
| [1-2H]-1,5,6-Tri-*O*-acetyl-2,3,4-tri-*O*-methylgalactitol | 6-Gal | 16.66 | 102 | 0.1 | 0.05 (50 % down) |
| [1-2H]-1,3,5,6-Tetra-*O*-acetyl-2,4-di-*O*-methylgalactitol | 3,6-Gal | 18.43 | 118 | 1.12 | 0.56 (50 % down) |
| [1-2H]-1,4,5-Tri-*O*-acetyl-2-methylacetamido-3,6-di-*O*-methylglucosaminitol | 4-GlcNAc | 20.86 | 117 | 1.08 | 0.93 (13 % down)[d] |
| [1-2H]-1,3,5-Tri-*O*-acetyl-2-methylacetamido-4,6-di-*O*-methylglucosaminitol | 3-GlcNAc | 21.57 | 117 | 0.65 | 0.18 (72 % down)[d] |

[a]RT – Retention time

[b]Area of most abundant EIC ion used for calculation

[C]The peak area was normalised to the peak area of non-reducing t-Man residue in each sample

[d]Quantification of HexNAc PMAA derivatives is less reliable than for hexose PMAAs.

### Table S2: List of oligonucleotide primers used in this study

| Oligo No. | | Sequence (5' – 3') | Description |
|---|---|---|---|
| ZJ1 | Forward | GGATCCAGGCCTCGGAGATCCTAACA | Amplification of pUC19 vector containing 5'UTR of Tb927.7.6170 and actin and 3'UTRs of actin and Tb927.7.6110 for Gibson assembly |
| ZJ2 | Reverse | AAGCTTCACTAGTTCTAGAGCTTATTTTATGGCAGCAA | |
| ZJ3 | Forward | CATAAAATAAGCTCTAGAACTAGTGAAGCTTATGGCCAAGCCTTTGTCTC | Amplification of blasticidin-S deaminase (BSD) drug resistance cassette for Gibson assembly |
| ZJ4 | Reverse | GTTAGGATCTCCGAGGCCTGGATCCTTAGCCCTCCCACACATAAC | |
| ZJ5 | Forward | GTAAAATTCACAAGCTTTAGATGGAGGTGGAACTGGAG | Amplification of Tb927.7.6170 for Gibson assembly (add back construct) |
| ZJ6 | Reverse | CCAACTAAATGGGCAGGATCcTTACACATGAATGCTCTTTCCCA | |
| ZJ7 | Forward | GTAAAATTCACAAGCTTTAGATGTACTCTGTTCATTGGGA | Amplification of Tb927.7.6150 for Gibson assembly (add back construct) |
| ZJ8 | Reverse | CCAACTAAATGGGCAGGATCcTTACACATGAATGCTCTTTCCCA | |
| ZJ9 | Forward | GTAAAATTCACAAGCTTTAGATGGAGGTGGAACTGGAG | Amplification of Tb927.7.6110 for Gibson assembly (add back construct) |
| ZJ10 | Reverse | CCAACTAAATGGGCAGGATCcTTACACATGAATGCTCTTTCCCA | |
| ZJ11 | Forward | GGATCCTGCCCATTTAGTTG | Amplification of plew100_v5 vector for Gibson assembly (add back construct) |
| ZJ12 | Reverse | CTAAAGCTTGTGAATTTTACTTTTTG | |
| ZJ13 | Forward | ATGACCGAGTACAAGCCCA | Amplification of PAC drug resistance cassette as Southern blotting probe |
| ZJ14 | Reverse | TCAGGCACCGGGCTTGCGGGTCA | |
| ZJ15 | Forward | ATGGCCAAGCCTTTGTCTC | Amplification of BSD drug resistance cassette as Southern blotting probe |
| ZJ16 | Reverse | TTAGCCCTCCCACACATAAC | |
| ZJ17 | Forward | ATGGAGGTGGAACTGGAGCCATTTG | Amplification of Tb927.7.6170 ORF as Southern blotting probe |
| ZJ18 | Reverse | TTACACATGAATGCTCTTTCCCA | |
| ZJ19 | Forward | GGTAAATTCGGTATTTCCCGCTG | Amplification of specific Tb927.7.6110 transcripts for RT-qPCR |
| ZJ20 | Reverse | ATTTCGAGTGCCGGGATCCG | |
| ZJ21 | Forward | CACCAAAGGGGCATACGTCA | Amplification of specific Tb927.7.6150 transcripts for RT-qPCR |
| ZJ22 | Reverse | AAAGCAATCACCACGGCAA | |
| ZJ23 | Forward | CGTAAACACTATTGTCCCAG | Amplification of specific Tb927.7.6170 transcripts for RT-qPCR |
| ZJ24 | Reverse | GCCATTACGAGGGCGAAA | |

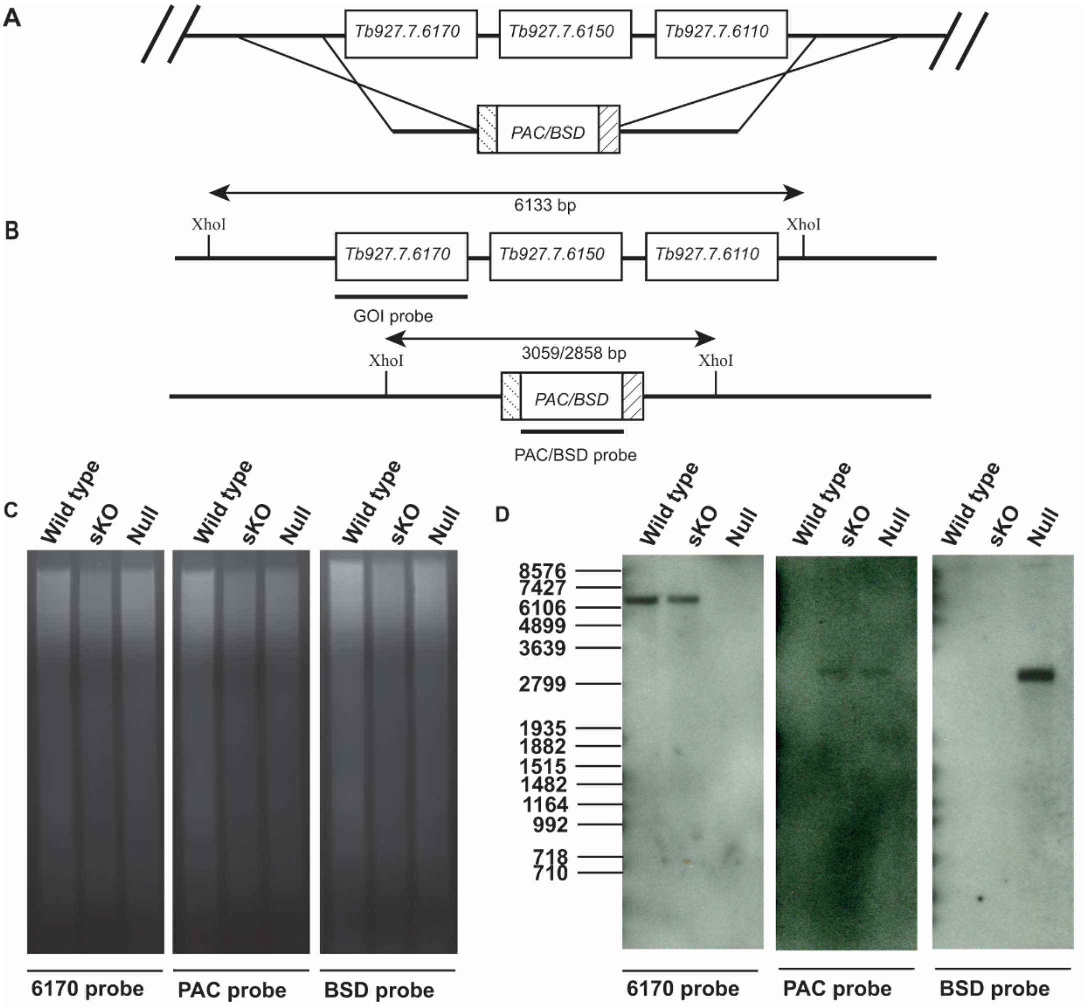


| | Expected band size (bp) | | |
|---|---|---|---|
| | GOI probe | PAC probe | BSD probe |
| Wild type | 6133 | No bands | No bands |
| sKO | 6133 | 3059 | No bands |
| Null $^{-/-}$ | No bands | 3059 | 2858 |

**Figure S1: Verification of the Tb927.7.6110/6150/6170$^{-/-}$ null mutant genotype by Southern blot.** (**A**) To create the Tb927.7.6110/6150/6170$^{-/-}$ null mutants, the first allele was targeted for replacement by *PAC* resistance cassette after which the second allele was replaced by *BSD* resistance cassette. (**B**) Schematic representation of predicted XhoI digestion sites and fragments to be detected by gene of interest (GOI), *PAC* and *BSD* probes for the Tb927.7.6110/6150/6170 locus. (**C**) and (**D**) Ethidium bromide staining and Southern blot, respectively of PCF *T. brucei* gDNA (5 µg/ lane) of wild-type and Tb927.7.6110/6150/617$^{+/-}$ single knock out (sKO) and Tb927.7.6110/6150/6170$^{-/-}$ null mutants digested with XhoI. The expected sizes of bands detected by the probes are indicated in (**B**) and in the table below the Southern blot.

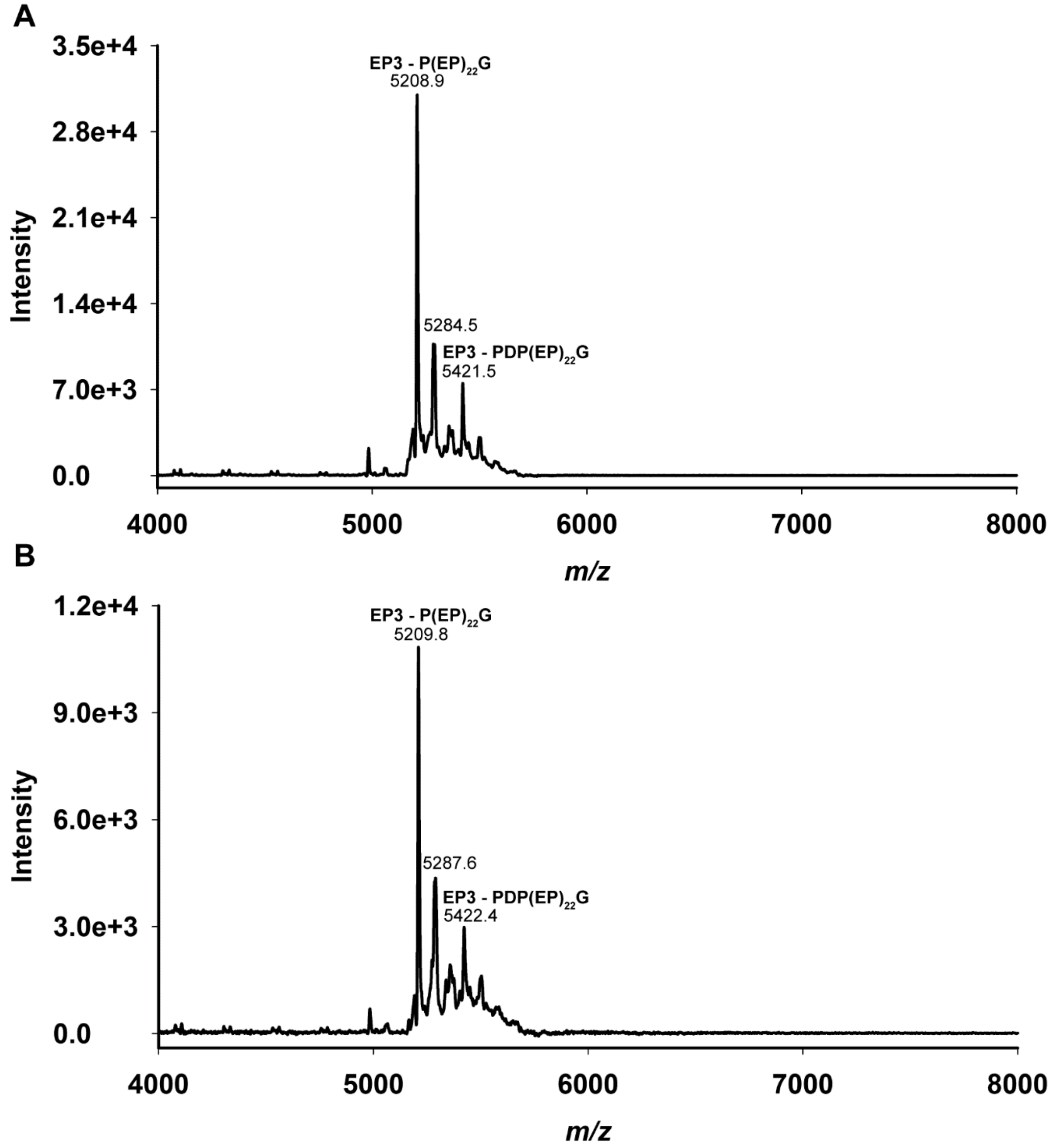


**Figure S2: Identification of procyclin types by MALDI-ToF** Procyclin samples from wild-type cells (**A**) and Tb927.7.6110/6150/6170$^{-/-}$ null mutant (**B**) were subjected to aqueous HF dephosphorylation and mild acid treatment (TFA) and analysed by negative-ion MALDI-ToF mass spectrometry (28). The species observed at *m/z* 5208 and 5421 represent EP3 procyclin fragments where ethanolamine is linked to the C-terminal glycine.

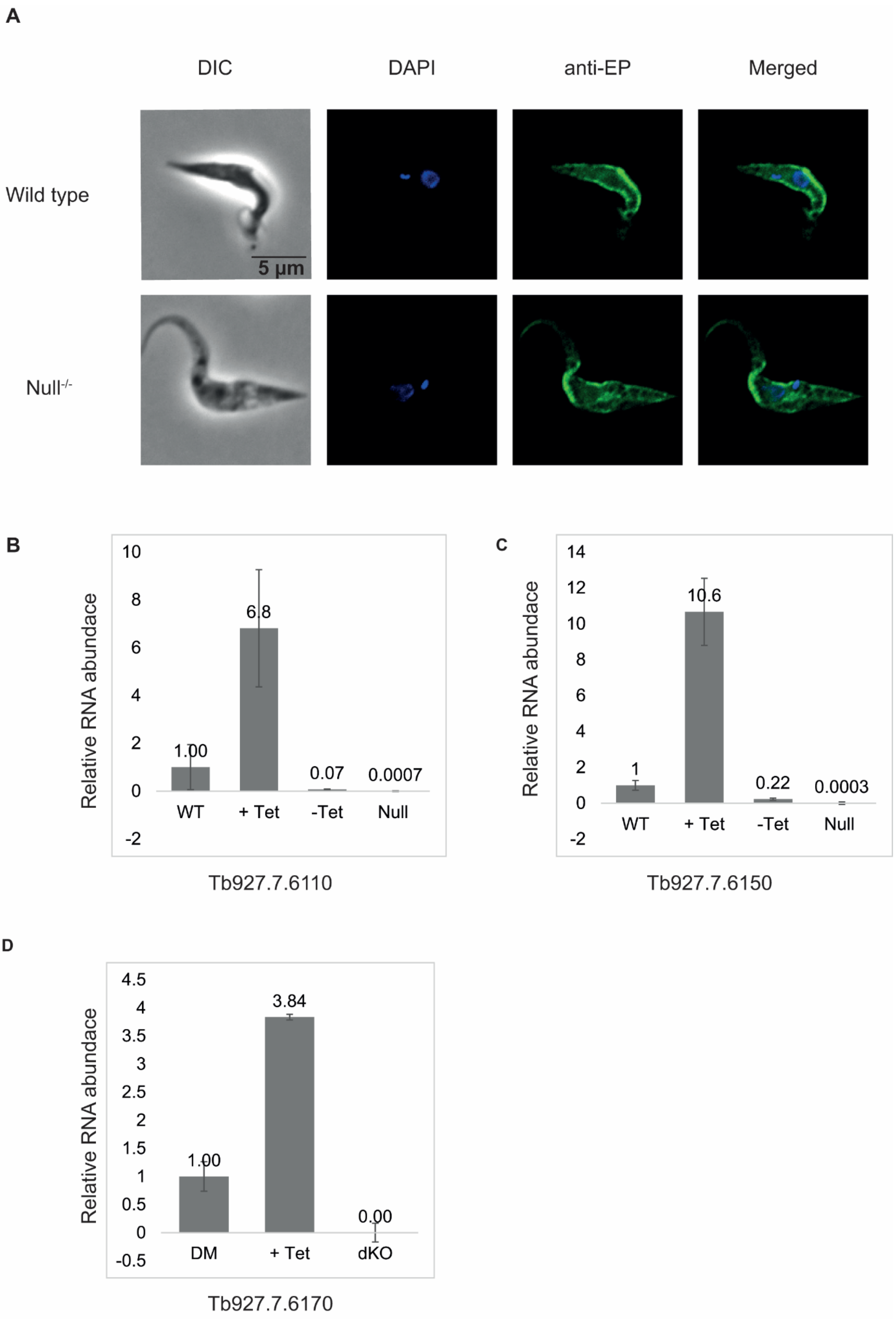

A
DIC
DAPI
anti-EP
Merged
Wild type
5 µm
Null-/-
B
Relative RNA abundace
10
8
6
4
2
0
-2
6.8
1.00
0.07
0.0007
WT
+ Tet
-Tet
Null
Tb927.7.6110
C
Relative RNA abundace
14
12
10
8
6
4
2
0
-2
10.6
1
0.22
0.0003
WT
+ Tet
-Tet
Null
Tb927.7.6150
D
Relative RNA abundace
4.5
4
3.5
3
2.5
2
1.5
1
0.5
0
-0.5
3.84
1.00
0.00
DM
+ Tet
dKO
Tb927.7.6170

**Figure S3: Immunofluorescence microscopy using anti-EP procyclin antibody in PCF wild-type and Tb927.7.6110/6150/6170$^{-/-}$ null mutants and RT-qPCR analysis of Tb927.7.6110, 6150 and 6170 transcript levels in wild-type, null and add back clones.**

(**A**) Fixed and permeabilized PCF wild-type *T. brucei* and Tb927.7.6110/6150/6170$^{-/-}$ null (Null$^{-/-}$) mutant parasites were stained with anti-EP procyclin antibodies to detect procyclins (green). The cells were also stained with DAPI to detect the nuclear and kinetoplast DNA and imaged by DIC. Similar cell surface staining patterns were observed in wild-type *T. brucei* and Tb927.7.6110/6150/6170$^{-/-}$ null mutant parasites.

(**B**), (**C**) and (**D**) Tb927.7.6110, 6150 and 6170 transcript levels in wild-type (WT) Tb927.7.6110/6150/6170$^{-/-}$ null mutant (Null) and Tb927.7.6110, 6150 and 6170 add back overexpressing cell lines, without (- Tet) and/or with (+ Tet) 24 h Tet induction, as determined by RT-qPCR. Wild-type served as normalisation control in each analysis, and the Tb927.7.6110/6150/6170$^{-/-}$ null mutant served as a negative control. The experiments were carried out in triplicate and the error bars indicate one standard deviation.

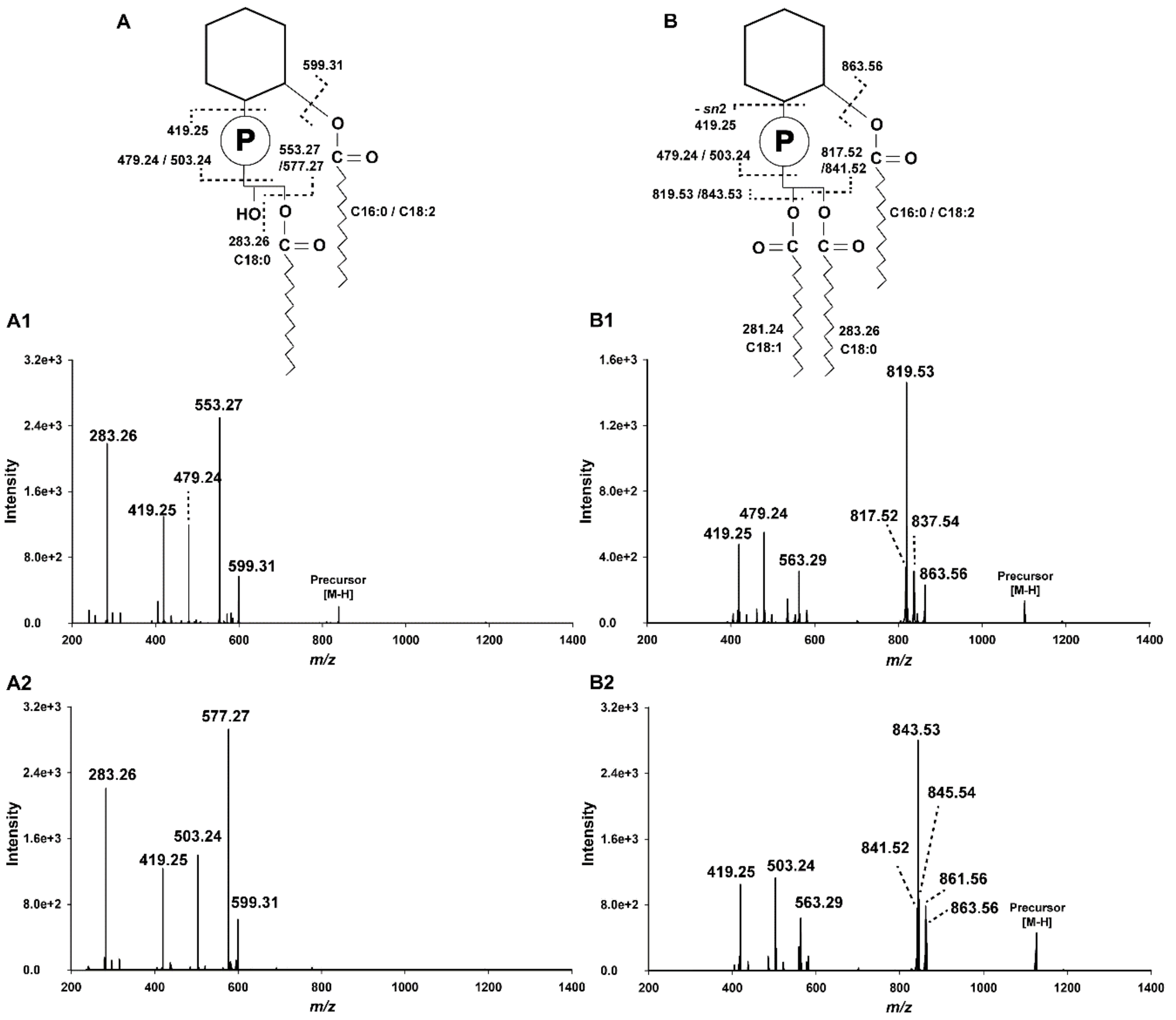


**Figure S4**: **ES-MS$^2$ identification of PI species**. The procyclin samples from wild-type and Tb927.7.6110/6150/6170$^{-/-}$ null mutant cells were subjected to nitrous acid deamination and the released PI species were analysed by ES-MS (Fig. 4) and ES-MS$^2$. (**A1**) and (**A2**) show the ES-MS$^2$ product ions of two major [M-H]$^-$ precursor ions observed in wild-type samples, m/z 837.55 and m/z 861.55, respectively. (**B1**) and (**B2**) show the ES-MS$^2$ product ions of two major [M-H]$^-$ precursor ions observed in Tb927.7.6110/6150/6170$^{-/-}$ null mutant samples, m/z 1101.80 and m/z 1125.80, respectively. The ES-MS$^2$ was fragmentation was acquired using collision induced dissociation. The product ion assignments for wild-type and Tb927.7.6110/6150/6170$^{-/-}$ null mutant PI species are indicated above in (**A**) and (**B**), respectively.
